# Sensing stress responses in potato with whole-plant redox imaging

**DOI:** 10.1101/2020.11.26.386573

**Authors:** Matanel Hipsch, Nardy Lampl, Einat Zelinger, Orel Barda, Shilo Rosenwasser

## Abstract

Environmental stresses are among the major factors that limit crop productivity and plant growth. Plant exposure to various abiotic stress, such as drought, cold temperatures, or high light, results in overproduction of reactive oxygen species (ROS). To avoid oxidative damage, critical mechanisms for their detoxification have evolved, consisting of ROS-scavenging enzymes and small antioxidant molecules, such as glutathione (GSH) and ascorbate. Thus, monitoring redox changes with high spatial and temporal resolution is critical for understanding oxidative stress signaling and has the potential to enable early detection of stress responses in crop plants. In this work, potato plants (*‘Solanum tuberosum’*) expressing a chloroplast-targeted reduction-oxidation-sensitive green fluorescent protein2 (roGFP2) were generated to report the redox potential of the glutathione (*E*_*GSH*_) in the chloroplast stroma. By applying whole-plant fluorescence imaging, we mapped alteration in the chloroplast *E*_*GSH*_ under several stress conditions including, high-light, cold and drought. Extremely high increase in chloroplast *E*_*GSH*_ was observed under the combination of high-light and low temperatures, conditions that specifically induce PSI photoinhibition. Intriguingly, whole-plant ratiometric imaging analysis noted a higher reduced state in newly developed as compared to mature leaves, suggesting a graded stress sensitivity as part of the plant strategies for coping with stress conditions. The presented observations suggest that whole-plant redox imaging can serve as a powerful tool for the basic understanding of plant stress responses as well as for applied agricultural research, such as improving phenotyping capabilities in breeding programs and early detection of stress responses in the field.

## Introduction

Crop plants live in highly dynamic environments, and abiotic stresses, such as salinity, drought, high temperature and high light, are thought to be the major constraints of crop production and ultimately of food security (Fahad et al., 2017). Continuous exposure to moderate stress levels or even to suboptimal growth conditions interrupts plant homeostasis, resulting in constant energy loss due to resource diversion towards the activation of defense and acclimation mechanisms (Zhu, 2016).

As a third major food crop, potato (*Solanum tuberosum)* productivity is crucial for worldwide food security (Devaux et al., 2014). For the past 20 years, there has been a dramatic increase in potato production and demand in Asia, Africa, and Latin America (da FAOSTAT, 2014). The potato tubers are rich sources of carbohydrates and provide essential nutrients, such as dietary fiber, vitamins, minerals, protein, and antioxidants (Bach et al., 2012). Despite, the wide distribution and adaptability of the potato plant to various environmental and climatic conditions, potato productivity is highly affected by environmental conditions (Bohnert, 2007). Even exposure to mild abiotic stresses such as drought and heat, as commonly occurs in potato growing regions, reduce photosynthesis efficiency, which significantly impacts potato production and quality (George et al., 2017). Thus, tools for sensing of early stress responses can provide important information regarding the complex signaling networks involved in potato stress acclimation strategies and has the potential to improve crop sustainability by enabling early intervention.

In the past few decades, there has been a growing demand for biosensing technologies that allow dynamic monitoring of crop growth and stress status, to support ‘on-line’ decisions ensuring the long-term sustainability of crop productivity. A range of technologies has been successfully implemented to nondestructively perceive information regarding the physiological status of plants, such as chlorophyll fluorescence imaging to detect photosynthetic activity (Wang et al., 2018), thermal imaging to estimate stomatal conductance and transpiration (Costa et al., 2013), multispectral imaging to evaluate crop water and nutrient status (Wang et al., 2018) and terahertz spectroscopy to detect plant drought stress responses (Born et al., 2014). However, technologies for early detection of the cellular biochemical signals involved in sensing and processing of environmental information regarding abiotic and biotic stress, are still lacking, mainly due to the destructive nature of common biochemical methods.

Stress-induced alterations in plant metabolism are typically accompanied by modifications in the levels of reactive oxygen species (ROS), which can differ in their chemical identity and subcellular localization (Foyer and Noctor, 2003; Gadjev et al., 2006; Miller et al., 2010; Møller and Sweetlove, 2010). Thus, increased ROS or oxidized metabolite levels, as well as the ROS-induced gene products, are common biomarkers for stress responses. Imaging and detection of ROS in live cells have been achieved using various fluorescent or dyes, such as 3,3’-diaminobenzidine (DAB), Amplex red, and 2’,7’-dichlorodihydrofluorescein diacetate (H_2_DCFDA) (Halliwell and Whiteman, 2004; Van Breusegem et al., 2008; Gao and Zhang, 2008; Zhang et al., 2009; Fichman et al., 2019; Fichman et al., 2020). A new type of nano-sensor probes, based on single-walled carbon nanotubes, was shown to enable spatiotemporal, real-time monitoring of H_2_O_2_ within plants (Lew et al., 2020; Wu et al., 2020). While these probes can be easily implemented in a wide range of plants, and provide valuable information on alterations in ROS metabolism on the whole-plant level they require the incorporation of the probes to specific plant tissues, making it challenging to investigate redox alterations on the whole-plant level over long physiologically relevant time periods.

Genetically encoded redox-sensitive green fluorescent protein (roGFP) probes have been developed and used as a biosensor in model systems. roGFP-based probes have allowed for *in vivo* monitoring of the glutathione redox potential (*E*_*GSH*_) and H_2_O_2_ at high spatiotemporal resolution (Dooley et al., 2004; Hanson et al., 2004; Jiang et al., 2006b; Meyer et al., 2007; Gutscher et al., 2008; Meyer and Dick, 2010; Nietzel et al., 2019). The roGFP mechanisms of action is based on two surface-exposed cysteine residues which form an intramolecular disulfide bridge that affects protein fluorescence. Due to their ratiometric nature, high sensitivity, reversibility, and insensitivity to pH alterations in the physiological range, roGFP-based redox sensors are powerful tools for investigating redox dynamics in subcellular compartments (Albrecht et al., 2011; Bratt et al., 2016; Ugalde et al., 2020). Their reversibility enables the monitoring of their redox state at multiple points over time without damaging the tissue. Notably, as the roGFP redox state is regulated by counteracting oxidative and reductive reactions, it reflects the transmission of redox signals, leading to modulation of the redox state of native redox-sensitive proteins as part of plant stress acclimation (Meyer, 2008; Rosenwasser et al., 2014). However, current approaches for measuring roGFP-based signals in plants are mainly based on confocal microscopy and plate readers, which are suitable for plant pieces or small model plants but not for intact crop plants grown in soil. The global importance of potato crop and the availability of highly efficient transformation protocols (Deblonde and Ledent, 2001; Eiasu et al., 2007; Schafleitner et al., 2007; Kumar et al., 2015) makes potato an ideal platform to investigate genetically encoded biosensor performance in crop plants.

In this work, potato plants expressing chloroplast-targeted roGFP2 were generated and subjected to whole-plant roGFP ratiometric imaging analysis using a highly sensitive *in vivo* imaging system. The experimental setup enabled detection of *in planta* redox modification in response to several abiotic stress conditions that mimicked natural field conditions. The presented data demonstrate that whole-plant imaging of roGFP-expressing plant allowed for spatially resolved mapping of stress-induced redox perturbations over long periods. These results may have important implications on phenotyping capacities in large-scale breeding projects and on the detection of stress conditions in crop plants under field conditions.

## Results and Discussion

### Generation of potato plants expressing genetically encoded redox probe

The roGFP probe allows for the quantitative, real-time readout of *E*_*GSH*_ in living cells (Meyer et al., 2007; Albrecht et al., 2011). To enable the monitoring of temporal alterations in the chloroplastic *E*_*GSH*_ (chl-*E*_*GSH*_) on the whole-plant level, *Agrobacterium*-mediated genetic transformation was applied to obtain potato plants cv. Desiree expressing the roGFP2 probe in the chloroplast (chl-roGFP2, Supp. Fig 1). Chloroplast targeting was achieved by using the *Arabidopsis* 2-Cys peroxiredoxin A signal peptide, which is targeted to the chloroplast stroma (König et al., 2002), as verified by the overlap of the chl-roGFP2 fluorescence with the chlorophyll fluorescence signals (Fig. 1A). No phenotypic differences between several independent lines expressing the roGFP2 probe and wild type were observed. The line with the highest fluorescence intensity and lowest variability in roGFP2 signals between individual plants was chosen for subsequent experiments. roGFP signal silencing, a phenomenon frequently reported for plant protein-based biosensors and has been observed to increase over generations, (Schwarzländer et al., 2016; Exposito-Rodriguez et al., 2017), was not detected, presumably since lines were propagated vegetatively either from cuttings or tubers.

**Figure 1:**
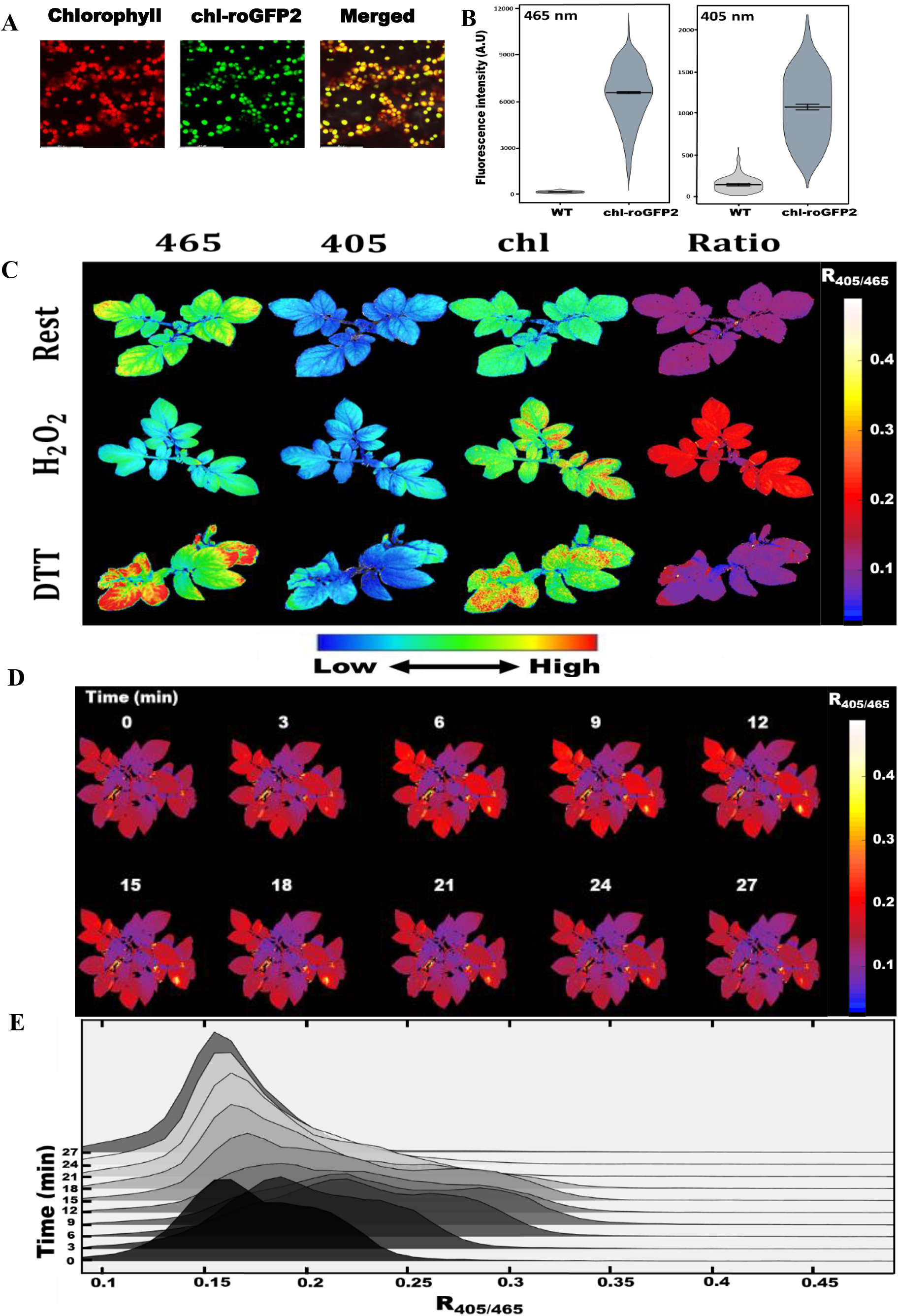
Live imaging of potato plants expressing roGFP2 in chloroplasts: A, Confocal images of chl-roGFP2-expressing potato plants showing its subcellular localization. B, Comparison between the emission intensities detected at 515nm in wild type (WT) and roGFP2-expressing plants, following excitation with 465 or 405nm. The data are summarized as a violin plot reflecting the pixel distribution of a representative plant. C, Fluorescence images and ratiometric analysis of chl-roGFP2 signals during rest and under fully oxidized (1000mM H2O2) and fully reduced (100mM DTT) conditions using whole-plant imaging. D, Whole-plant timeline ratiometric images of potato plant watered with 1M H2O2 and monitored. The numbers on the top represent the time from H2O2 application. E, Ridgeline plot showing the pixel distributions of ratio images of a representative plant.

### *In vivo* redox imaging of potato biosensor plants

The roGFP2 probe emission at 515 nm was recorded using a highly sensitive *in vivo* imaging system, following excitation at 405 and 465nm with light-emitting diodes (LEDs). Ratio values (R_405/465_) obtained from the division of the emitted fluorescence due to excitation at 405◻nm versus 465◻nm indicated the roGFP2 oxidation state (Dooley et al., 2004; Hanson et al., 2004). To explore the ability of this set-up to provide reliable information on *in planta* chloroplastic specific redox alterations on a whole-plant level, chl-roGFP2 fluorescence was monitored in intact plants grown in soil. To estimate the plant auto-fluorescence level, the fluorescent signal obtained for plants expressing the roGFP2 probe was compared to the signal measured in wild type plants. As shown in Figure 1B, a clear separation between the pixel intensity histograms of roGFP2-expressing versus wild type lines was observed following excitation at 465 nm and 405 nm, demonstrating an intense roGFP2 signal in the former lines. The measured autofluorescence signals were 2.2% and 9.6% of the total fluorescence signals following excitation at 465nm and 405nm, respectively. In all subsequent experiments, autofluorescence values obtained from wild-type plants exposed to the same conditions as roGFP-expressing lines were used for background correction (See Methods). While in chl-roGFP2-expressing plants autofluorescence values were relatively minor, such correction is highly significant for biosensor lines exhibiting less intense fluorescence signals, as commonly observed in plant lines expressing fluorescent probes in other cellular components (e.g., mitochondria and peroxisomes) and roGFP2 fusion lines. Correction for background values is also highly important under specific stress conditions that result in increase autofluorescence, especially in the 405nm excitation range (Rosenwasser et al., 2010).

To monitor the *in planta* response to redox changes, 2.5-week-old chl-roGFP2-expressing plants grown from cuttings, were imaged under steady-state conditions and following the application of H_2_O_2_ and DTT. While some variability in absolute fluorescence roGFP intensities were observed among different leaves, the ratiometric analysis obtained by division of the 405 nm image by the 465 nm image, pixel by pixel, resulted in informative false-color images (Fig. 1C). Upon H_2_O_2_ treatment, an increase in 405 nm-excited fluorescence and a decrease in 465 nm-excited fluorescence were observed, demonstrating probe oxidation. Conversely, the treatment of plants with DTT resulted in probe reduction, as demonstrated by the less intense signal following 405 nm excitation and increased brightness following 465 nm excitation. Ratiometric images of untreated plants demonstrated the highly reduced chl-*E*_*GSH*_ throughout the whole plant.

The temporal response of 4-5-weeks-old plants to oxidative conditions was then assessed by watering of soil-grown chl-roGFP2-expressing plants with 50ml 1M H_2_O_2_. Ratiometric images and the derived pixel distribution analysis demonstrated an increase in the chl-roGFP2 oxidation state, which became saturated after 12 min of exposure, and was then followed by reduction, reaching a steady-state level after 27 min (Fig. 1 D&E, Supp Movie 1). Interestingly, newly developed leaves on the higher part of the plant exhibited a higher reduced state of chl-roGFP2 as compared to lower and mature leaves under steady-state. These spatial differences were also observed following the H_2_O_2_ application and can result from a differential antioxidant activity of young and old leaves or unequal distribution of H_2_O_2_ among the plant tissue (Fig. 1 D&E, Supp Movie 1). These results showed quantitative *in vivo* mapping of the chl-roGFP2 oxidation state and demonstrated its kinetics on a whole-plant level.

To further calibrate the probe dynamic range, detached leaves were soaked in various H_2_O_2_ concentrations and the redox state was monitored. Treatment of leaves with increasing H_2_O_2_ concentrations (10mM to 1500mM) resulted in increased R_405/465_, as visualized by ratiometric images and pixel ratio distributions (Fig. 2, Supp. Fig 2). Kinetic measurements demonstrated that chl-roGFP2 fluorescence reached saturation (R_405/465_=0.49) after 12 min, at relatively high H_2_O_2_ concentrations of 750-1000mM (Fig. 2A). Notably, these values do not reflect the endogenous H_2_O_2_ concentrations, which are affected by penetration rates and detoxification activity. Treatment of plants with 100mM DTT resulted in a decrease in R_405/465_ (0.09), demonstrating a dynamic range (R_405/465_ under oxidized conditions divided by R_405/465_ under reduced conditions) of 5.4, which is comparable to values obtained using confocal microscopy and a plate reader (Schwarzländer et al., 2008; Rosenwasser et al., 2010). The oxidation degree (OxD) of chl-roGFP2 and chl-*E*_*GSH*_ in whole plants under steady-state conditions was calculated using the reference ratio values for fully oxidized and reduced states according to (Meyer et al., 2007). The average OxD values were approximately 25%. Considering a stromal pH of 8, this OxD value would reflect an *E*_*GSH*_= −346mV, which aligns with previous calculations of chloroplastic *E*_*GSH*_ under steady-state conditions in *Arabidopsis* plants (Schwarzländer et al., 2008; Rosenwasser et al., 2010). Interestingly, newly developed and mature leaves exhibited an average chl-roGFP2 OxD of 14% and 28%, respectively, demonstrating a deviation of 11 mV between different leaves on the same plant (Supp. Fig. 3).

**Figure 2:**
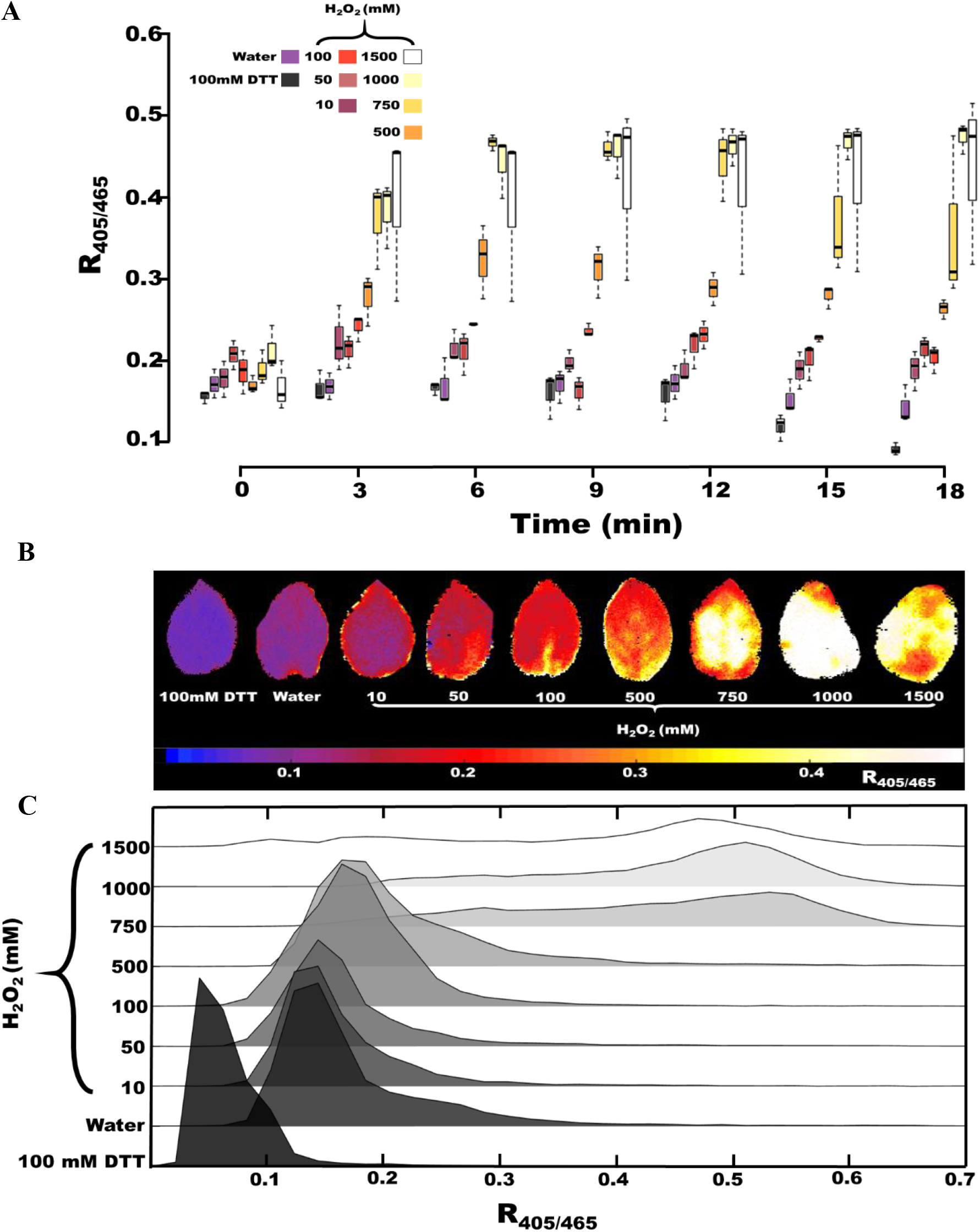
Redox sensitivity of the chl-roGFP2 probe to oxidative conditions in potato leaves. A, Does-dependent and temporal response of chl-roGFP2 to increasing concentrations of H2O2, as evaluated using ratiometric fluorescence imaging. R405/465 values are presented as a box plot (n=3) B, C Ratiometric images and pixel distribution analysis of a representative leaves 12 min after H2O2 application or 60 min after application of DTT.

### Daily measurements of chl-*E*_*GSH*_ under high light

Potato biosensor plants offer the opportunity to examine the *in vivo* influence of environmental stress conditions on the chl-*E*_*GSH*_ and to examine possible redox differences throughout plants. As the chl-*E*_*GSH*_ dynamically responds to changes in light intensities (Haber and Rosenwasser, 2020), we sought to apply the chl-roGFP2 probe to investigate the response of potato plants to changes in light intensities that mimic the kinetics and light intensities of field conditions. To this end, four-week-old plants were exposed to the following 16 h light conditions: Constant light (CL)-200 μmol photons m^−2^ s^−1^, Medium-light (ML) - light intensity was gradually increased, reaching a maximum value of 720 μmol photons m^−2^ s^−1^, followed by a gradual decrease toward the end of the day, High-light (HL) – gradual increases in light intensity, up to a maximum value of 1250 μmol photons m^−2^ s^−1^, followed by a gradual decrease (Fig. 3A). RoGFP fluorescence images were taken every two hours, starting from the light onset. As shown in Figure 3, the exposure of plants to increasing light intensities (ML and HL) resulted in an increased whole-plant chl-roGFP2 oxidation state, reaching maximum values in the middle of the day. Similar oxidation dynamics were observed in plants exposed to ML and HL, with slightly higher oxidation values under the HL treatment. For example, at 8 h from light onset, OxD values of 43±6% and 47±6%, were recorded in plants exposed to ML and HL, respectively. Relatively constant chl-roGFP2 OxD, of approximately 30%, was measured throughout the day in plants exposed to CL. Such a deviation between HL compared to CL in the roGFP2 oxidation state is equivalent to a 10mV increase in chl-*E*_*GSH*_, in agreement with values obtained in *Arabidopsis* plants under high light conditions (Haber and Rosenwasser, 2020). Returning to steady-state values was detected 14 h after light onset, when light intensity was reduced to 200 μmol photons m^−2^ s^−1^. The inspection of ratiometric images revealed that although a light-induced oxidation response was detected in all plant leaves, the differences between the chl-roGFP2 oxidation state in newly developed versus mature leaves were maintained under HL conditions. (Fig 3B). Taken together, these observations demonstrate that spatial heterogeneity and physiological responses of chl-*E*_*GSH*_ to light intensity can be nondestructively monitored in crop plants grown in soil. As high light induces H_2_O_2_ production (Exposito-Rodriguez et al., 2017), the increase in chl-*E*_*GSH*_ is likely a reflection of a new balance point between photosynthesis-dependent ROS production and NADPH-dependent glutathione reductase (GR) activity. As roGFP2 oxidation-reduction dynamics may reflect similar patterns occurring in many redox-regulated metabolic proteins, specifically those regulated by native GRXs (Meyer, 2008; Rosenwasser et al., 2014), it may provide an important readout of stress-induced metabolic alterations.

**Figure 3:**
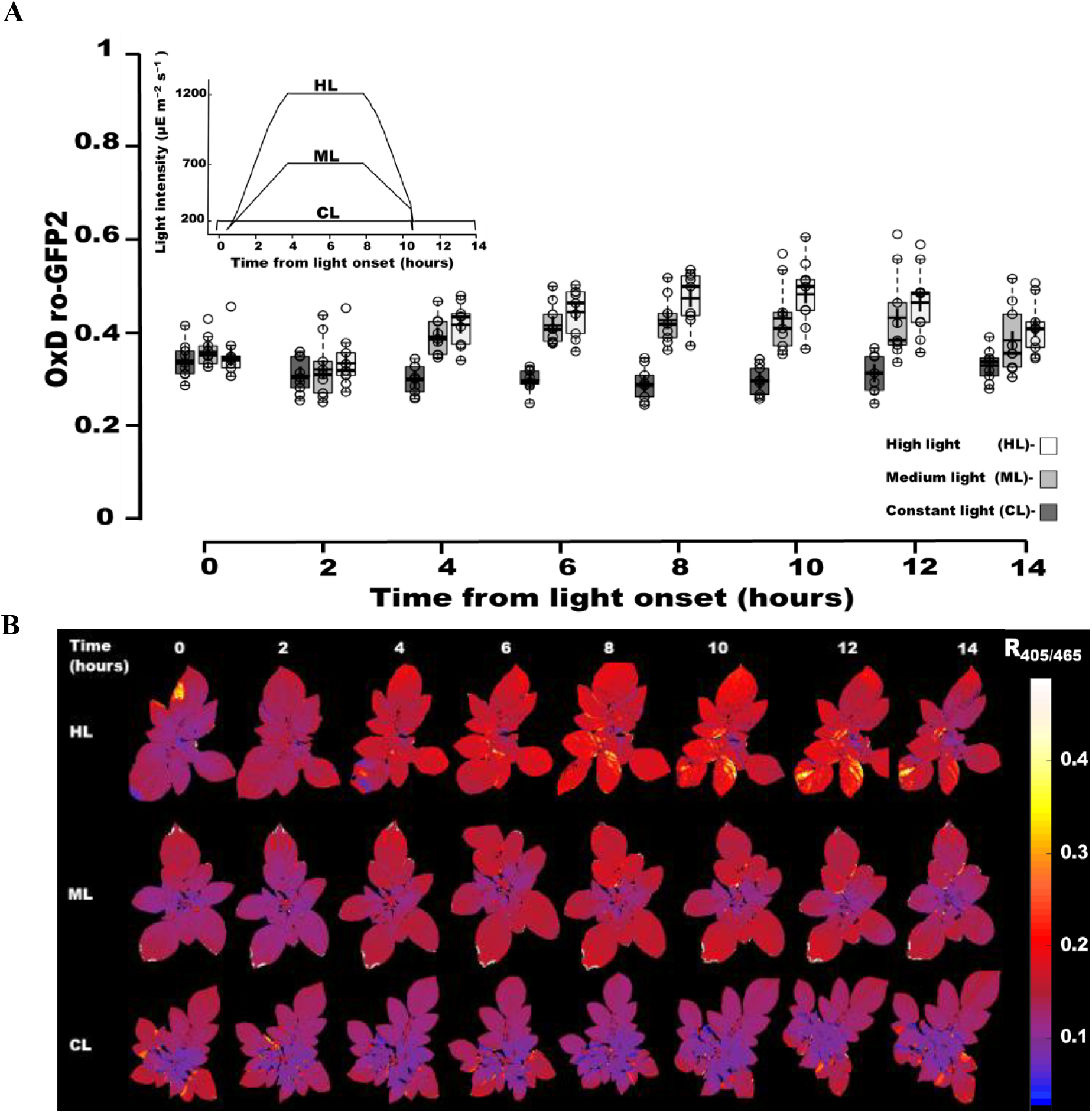
Daily imaging of the redox state of chl-roGFP2 under high light. A, Changes in chl-roGFP2 degree of oxidation (OxD) in response to changes in light intensity. Data are presented by box plots (n=9). Inset: daily light intensity regimes. B, Whole-plant ratiometric images of representative potato plants at different time points during the exposure to the light regimes presented in A. The color code highlights the spatial distribution of redox changes. The numbers on the top represent the time (h) from light onset.

### Profound chl-*E*_*GSH*_ oxidation under the integration of high light and cold temperatures

Various environmental conditions lead to an imbalance between photosynthesis light absorption and downstream carbon assimilation reactions, resulting in increased plant sensitivity to excess light energy. Specifically, photosystem I (PSI) photo-inactivation was observed in chilled potato leaves exposed to high light, likely due to ROS accumulation (Havaux and Davaud, 1994). Such stress combinations are of interest, as they result in yield reduction in crops and may pose a risk to plant tissues. To examine the effect of chilling stress on the chl-*E*_*GSH*_, chl-roGFP2-expressing plants were exposed to a low temperature (3°C) and to each of the three daily light treatments mentioned above (Fig. 3); whole-plant oxidation patterns were measured every two hours during the light period. In plants exposed to 3°C + CL, chl-roGFP2 OxD reached a stable state of approximately 50%, with a slight reduction after 2 h in the light period (Figure 4A). In contrast, marked oxidation levels were observed in plants exposed to the combination of cold temperature and high light, reaching chl-roGFP2 OxD values of 60% and 80% in plants exposed to 3°C+ ML and HL, respectively, after 10 h in the light. No decrease in OxD levels in 3°C+ ML or 3°C+ HL was observed when the light dimmed to lower light intensities at the end of the day. Remarkably, newly developed leaves on the upper part of the plants, particularly near the meristem, remained highly reduced, throughout the day, even under these harsh conditions, demonstrating heterogeneous responses between old and young leaves. Taken together, these results indicate that low temperature and high light conditions induced extensive chl-*E*_*GSH*_ oxidation, reaching levels comparable to those measured after application of extremely high concentrations of exogenous H_2_O_2_, implying a correlation between PSI photoinhibition and chl-*E*_*GSH*_ oxidation.

**Figure 4:**
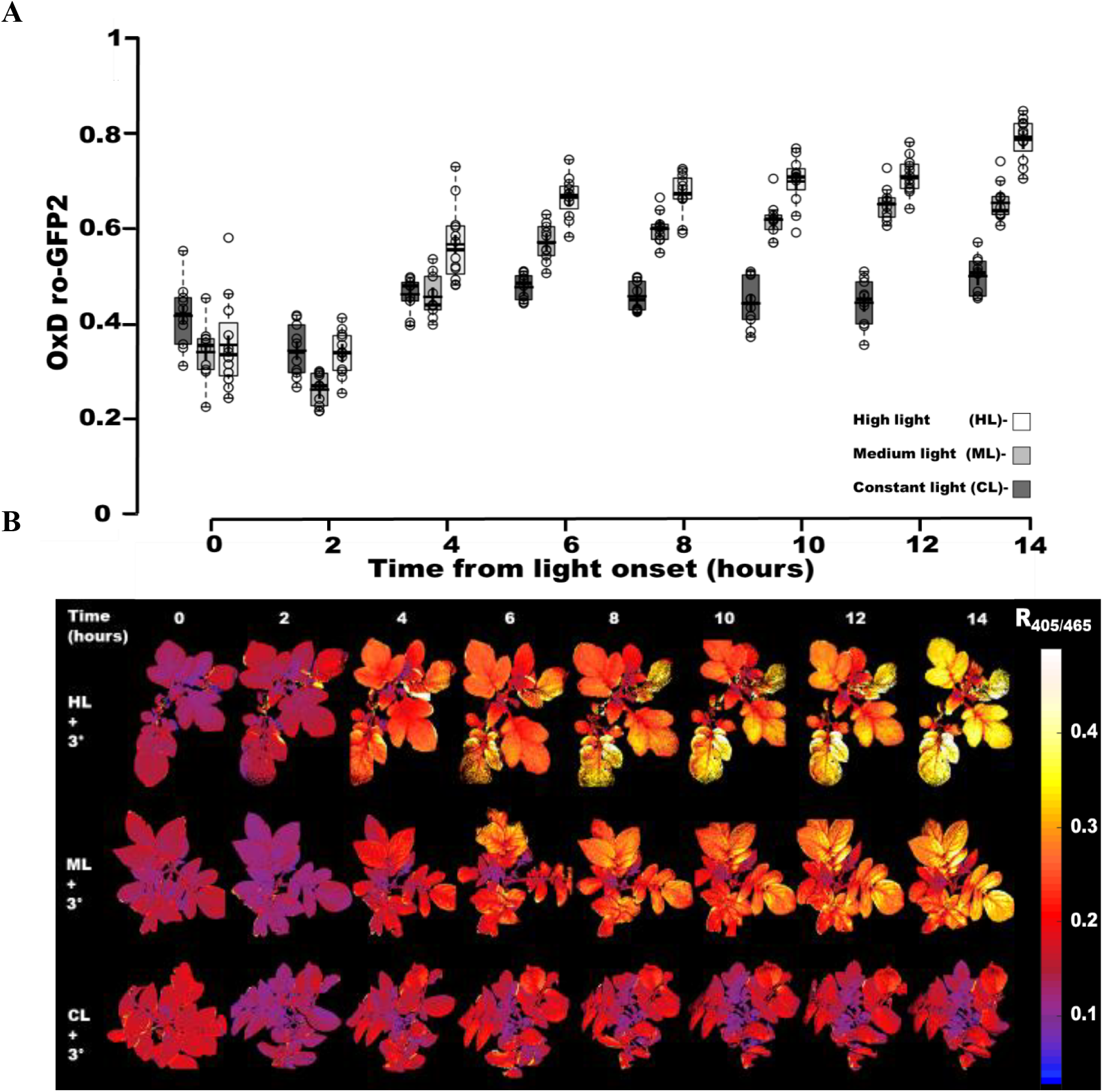
Monitoring of the redox state of chl-roGFP2 under combined high light and cold stress. Plants were exposed to cold temperatures (3°C) under the same light regime as in Fig. 3. A box plot presentation of the changes in chl-roGFP2 degree of oxidation (OxD) (n=10-12). B, Whole-plant timeline images of representative potato plants, highlighting the spatial distribution of redox changes. The numbers on the top represent the time (h) from light onset

The demand for reduced GSH under chilling stress was also demonstrated in transgenic tomato plants with suppressed glutathione reductase activity, and correlated with aggravated PSI photoinhibition and delayed PSI recovery (Shu et al., 2011). The extremely high roGFP2 oxidation state and the fact that no reduction was observed after returning plants to normal light conditions, suggests that plants failed to acclimate to such intensive stress. The significant changes in chl-*E*_*GS*__H_ (~32 mV) may indicate the induction of cell death as a similar range of changes in the chloroplast and mitochondrial *E*_*GS*__H_ was associated with cell death in *Arabidopsis* and diatoms (Rosenwasser et al., 2014; Van Creveld et al., 2015; Bratt et al., 2016; Volpert et al., 2018).

Higher activity of several photoprotective mechanisms, including upregulation of energy dissipation via heat and photorespiration, in young compared to mature leaves was observed in crop plants (Bertamini and Nedunchezhian, 2003; Jiang et al., 2006). The higher reduced state of the chl-*E*_*GSH*_ in young leaves may result from their decreased photosynthetic activity, raising the threshold at which light causes increased ROS production and subsequent chl-*E*_*GSH*_ oxidation. While the induction of these photoprotective pathways results in suboptimal photosynthetic efficiency, it may protect the photosynthetic machinery from severe destruction. Thus, the co-existence of leaves with differential chl-*E*_*GSH*_ values can be viewed as an evolutionary compromise between photosynthetic efficiency and photo-protection, enabling young leaves to better withstand unpredicted increases in light intensities and older leaves to photosynthesize more efficiently.

### Drought stress initiates early oxidation of chl-*E*_*GSH*_

Stomata closure in response to water stress restricts CO_2_ diffusion and reduces Calvin-Benson Cycle reactions, resulting in higher ROS production due to channeling of excessive light energy toward molecular oxygen (Suzuki et al., 2012). An increase in GR activity was reported under HL and water stress conditions, suggesting the increased activity of the ascorbate-GSH cycle (Yang et al., 2008; Gill and Tuteja, 2010). To examine whether drought-induced redox alterations in chloroplast can be monitored *in vivo*, chl-roGFP2 oxidation dynamics were followed during drought stress. Watering of four-week-old plants was stopped, and roGFP imaging was acquired every day (from the third day of irrigation stop). Starting six days after water was withheld, a gradual increase in chl-roGFP2 OxD levels was noted, with oxidation initiating in the peripheral leaves, and later spreading to all plant leaves (Fig. 5A&D). These observations are in line with the differential activation of the antioxidant response and energy dissipation pathways in young versus mature *Arabidopsis* leaves under drought conditions (Jung, 2004; Sperdouli and Moustakas, 2014).

**Figure 5.**
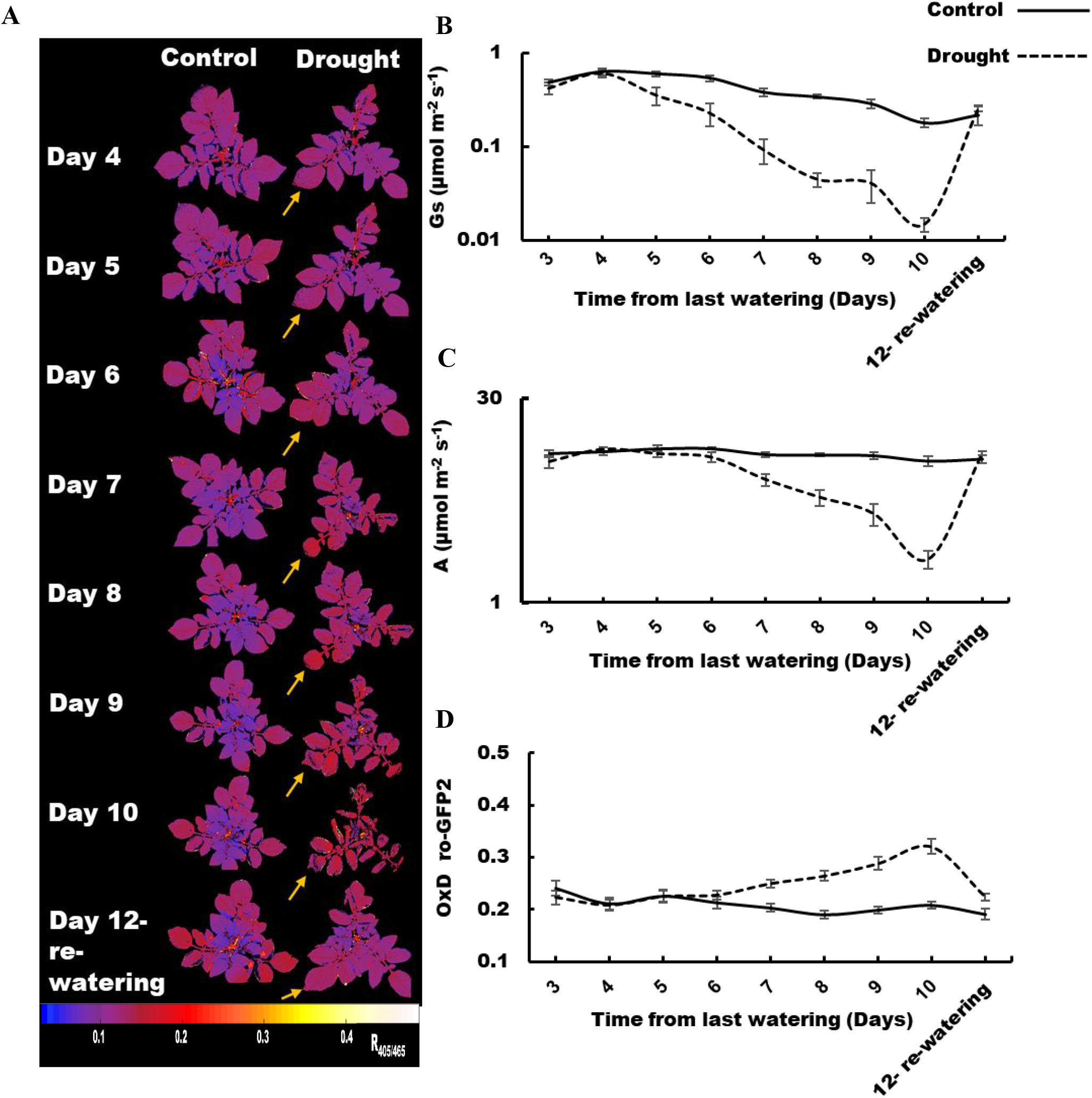
Changes in the redox state of chl-roGFP2 in well-watered and water-stressed plants. Drought conditions were induced by discontinuing irrigation. A, Whole-plant timeline images of representative well-watered and water-stressed potato plants. The yellow arrows indicate the oxidation changes in sample leaves during the course of drought stress. The numbers on the top represent the day from irrigation termination. B) Stomatal conductance (GS). C, Assimilation rate (A) D, Oxidation degree of chl-roGFP2 (OxD).

The rise in chl-roGFP2 OxD perfectly paralleled the decrease in carbon assimilation rates (Fig. 5C), pointing to the consistent relationship between photosynthetic activity and chl-*E*_*GSH*_ and suggesting that the increase in chl-*E*_*GSH*_ reflects an imbalance between light absorption and the ability to utilize it in the Calvin-Benson Cycle. The decline in stomatal conductance observed one day before the increase in chl-roGFP2 OxD (Fig 5B) further supports the notion that a decrease in intracellular CO_2_ levels resulted in an increase chl-*E*_*GSH*_. Steady-state redox levels and photosynthetic parameters were restored on day 12, upon re-watering of water-stressed plants, demonstrating the ability to sense the reversibility of stress response by the used probe. The observed increase in chl-*E*_*GSH*_ in water-stressed plants may reflect a broader increase in cellular ROS production, as suggested by the oxidation of the cytosolic and mitochondrial *E*_*GSH*_ reported for Arabidopsis plants (Jubany-Mari et al., 2010; Bratt et al., 2016). Future evaluations of organelle-specific changes in *E*_*GSH*_ using plant lines expressing the roGFP2 probe in various subcellular compartments will enable dissection of the exact microenvironment in which redox alterations are initiated under water stress conditions.

The GSH redox state is widely used as a marker of oxidative stress (Noctor et al., 1998; Schafer and Buettner, 2001; Dietz, 2003; Kranner et al., 2006; Meyer, 2008). Specifically, monitoring redox changes in photosynthesizing chloroplasts can provide valuable data regarding the response of leaf photosynthesis to environmental stresses. This is of great interest in crop plants in which plant productivity is greatly affected by the level of stress imposed on the photosynthetic machinery. The presented data suggest that crop plants expressing genetically encoded fluorescent sensors that report the chloroplast *E*_*GSH*_ can be a powerful tool to evaluate plant stress responses. Its applications can be broad, including, for example, in testing the performance of various chemicals in improving plant tolerance under stress conditions. The generated reporterplants can also be applied to improve high-throughput plant phenotyping in plant breeding programs and ultimately serve as highly sensitive tools for early detection of stress responses in the field. Notably, the need for highly sensitive fluorescence cameras may limit the accessibility to this technology, raising the need to develop portable and sensitive instruments to monitor GFP-based signals in the field. Taken together, given the significant role of redox metabolism in plant acclimation to stress conditions, crop plants expressing redox sensors can expand the basic understanding of plant stress physiology and extend the arsenal of early noninvasive tools for detection of stress-induced physiological changes in crops.

## Materials and Methods

### Plant material, growth conditions, and experimental set-up

Wild type and chl-roGFP2-expressing potato plants were planted in moist soil in 26.82 x 53.49 cm pots and placed in a controlled-environment greenhouse. Plants were vegetatively propagated from cuttings. All experiments were performed on 3-4-week-old plants in a FytoScope FS-RI 1600 plant growth chamber (Photon Systems Instruments). Plants were moved to the chamber several days before the experiments to allow acclimation to the chamber environment. In all experiments, plants were incubated in 60-70% relative humidity (RH) and ambient CO_2_.

### Production of transgenic plants

Potato leaves (cv. Désirée) were used for *Agrobacterium*LJmediated infection as described previously by (Ooms et al., 1987; Teper◻Bamnolker et al., 2017). Leaves were taken from clean culture and infected by *Agrobacterium tumefaciens* strain LBA 4404 harboring the pART27 plasmid, which contains the chl-roGFP2 construct (Haber and Rosenwasser, 2020). After the inoculation, cultures were transferred into cali induction medium containing kanamycin 50 (mg/L), cefotaxime 500 (mg/L), 6-Benzylaminopurine (BAP) 0.1 mg/L and 1-Naphthaleneacetic acid (NAA) 5mg/L, for 10 days. Then, plants were transferred to shot induction medium (regeneration media) containing Zeatin-riboside (ZR) 2mg/L, NAA 0.02 mg/L Gibberellic acid 3 GA3 0.02mg/L and appropriate antibiotics. Medium was refreshed every 14 days until shoots appeared. Transgenic explants from cultures that were fully regenerated with no roots were transferred to moist soil for rooting and sorting. Appropriate lines were selected by evaluating the chl-roGFP2 fluorescence signal (Supp. Fig 1).

### Confocal microscopy

Images were acquired with a Leica TCS SP8 confocal system (Leica Microsystems) and the LAS X Life Science Software, while using a HC PL APO ×40/1.10 objective. All images were acquired at a 4096×4096-pixel resolution, with emission at 500-520 nm following excitation at 488nm for chl-roGFP2 fluorescence and emission at 670nm following excitation at 488nm for chlorophyll fluorescence. Merged images were generated using Fiji (Image 1.A) software.

### Gas exchange measurements

Measurements of carbon assimilation rates were made using the portable gas analyzer Li-Cor-6800 gas exchange (LICOR, Lincoln, NE, USA). Environmental conditions in the leaf chamber were defined as a constant CO_2_ level of 400 ppm. Light intensities were similar to those applied in the growing chamber. The temperature and humidity were set at 25°C and 70%. Net photosynthesis (A) and stomatal conductance to water vapor (Gs) were measured. Between 6-9 plants were randomly selected for each treatment and measurements were taken at 12 PM each day.

### Chl-roGFP2 fluorescence measurements and image analysis

Whole-plant chl-roGFP2 fluorescence was detected using an Advanced Molecular Imager HT (Spectral Ami-HT, Spectral Instruments Imaging, LLC., USA), and images were acquired using the AMIview software. For chl-roGFP2 fluorescence detection, excitation was performed with 405nm±10 or 465nm±10 LED light sources and a 515nm±10 emission filter was used. For chlorophyll detection, a 405nm±10 LED light source and 670nm±10 emission filter were used. All images were taken under the same settings of LED intensities and exposure time. Chlorophyll autofluorescence was measured to generate a chlorophyll mask, which was then used to select pixels that returned a positive chlorophyll fluorescence signal. Only those pixels were subsequently considered for the roGFP analysis. For background correction, the average signal of WT plants without chl-roGFP2 was determined and subtracted from the values detected in the chl-roGFP2 fluorescence analysis. Ratiometric images were created by dividing, pixel by pixel, the 405 nm image by the 465 nm image, and displaying the result in false colors. Images were preprocessed using a custom-written Matlab script.

For calibration of the probe response, detached fully expanded leaves were immersed in 1M H_2_O_2_ and 100 mM DTT, and ratiometric images for fully oxidized and fully reduced states, respectively, were then acquired. RoGFP2 OxD (The relative quantity of oxidized roGFP proteins) was calculated for individual plants based on the whole-plant fluorescence signal according to Equation 1(Meyer et al., 2007).

Equation 1:

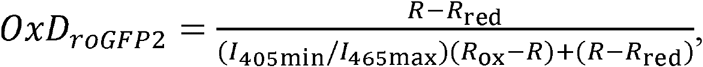

where R represent the 405/465 fluorescence ratio at the indicated time and treatment, R_red_ represents the 405/465 fluorescence ratio under fully reduced conditions, R_ox_-represents the 405/465 fluorescence ratio under fully oxidized conditions, I465_ox_ -the fluorescence emitted at 515 nm when excited at 465 nm under fully oxidized conditions and I480_red_ -the fluorescence emitted at 515nm when excited at 465nm under fully reduced conditions. E_GSH_ values were calculated according to Schwarzlander et al. (2008).

## Author contributions

M.H. and S.R. conceived and designed the research. M.H., N.L., E.L., and O.B. performed the research. M.H. and S.R. analyzed the data. M.H. and S.R. wrote the manuscript.

## Acknowledgments

We thank Andreas J. Meyer for his critical comments on the manuscript. We thank Dani Eshel and Paula Teper-Bemnolker for providing the starting materials and protocol for potato transformation. This research was supported by the Israel Science Foundation (grant No. 826/17 and No. 827/17) to SR.

