## Supplementary Figures for "Sensing stress responses in potato with whole-plant redox imaging"

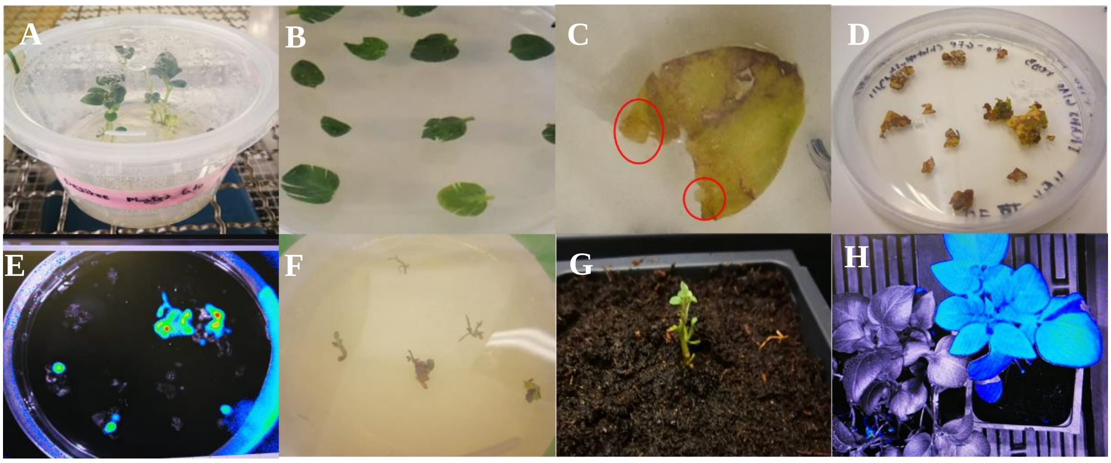


**Supplementary Figure 1**: Schematic steps of *Solanum tuberosum* 'Desiree' transformation. A, WT clean culture. B, Leaves infected by Agrobacterium. C, Callus development. D, Mature calli development. E, Screening of transgenic calli expressing roGFP by in vivo imaging system. F, Callus undergoes full dedifferentiation. G, Explant moved to the soil for rotting. H, Whole plant imaging of WT (left side) and transgenic plant (right side) expressing roGFP2 in the chloroplast.

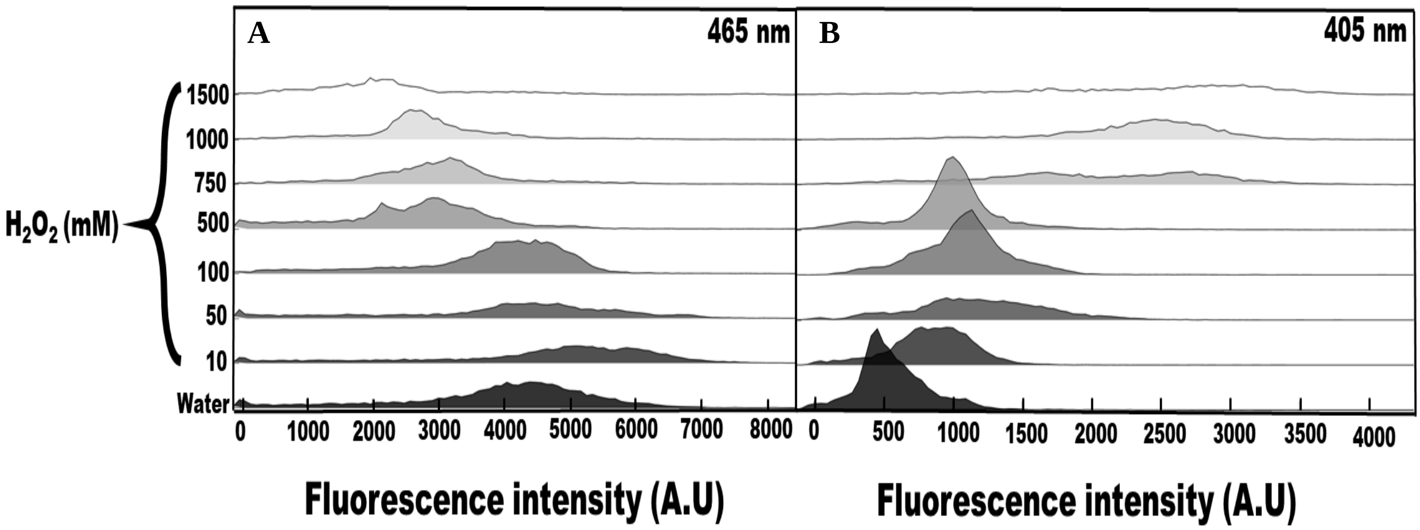


**Supplementary Figure 2:** Ridgeline plot showing pixel distributions of the emission intensity at 515nm of representative leaves following excitation with 465nm (A) or 405nm (B), in response to increasing $H_2O_2$ concentrations.

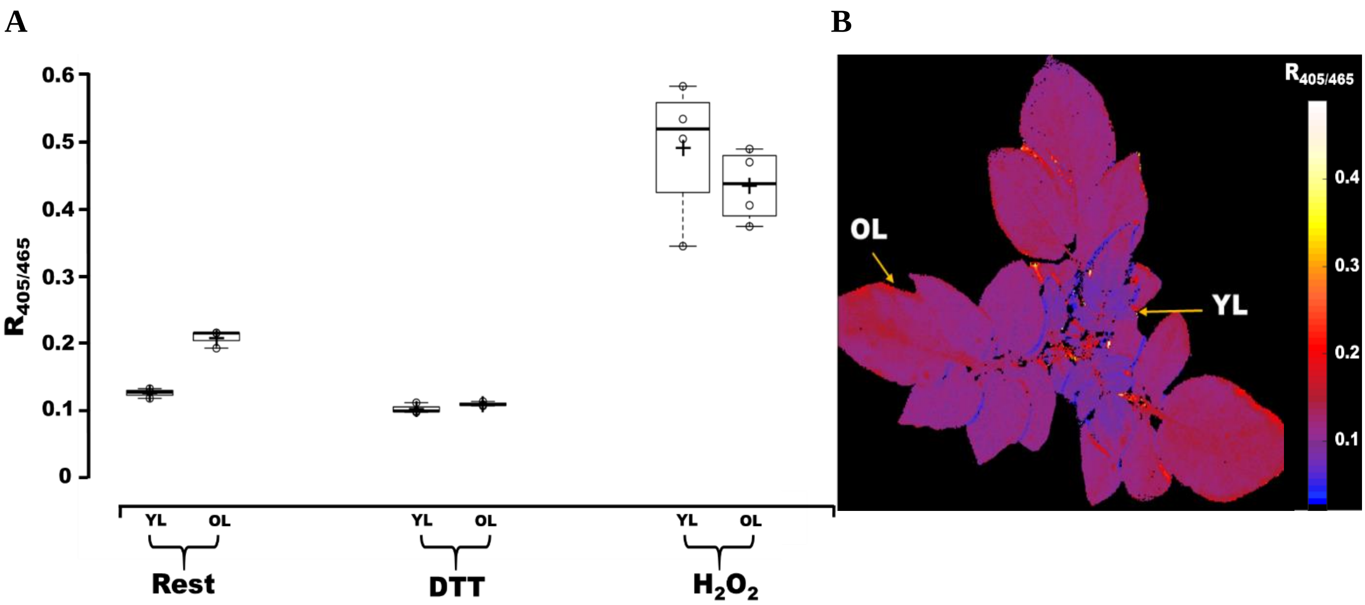


**Supplementary Figure 3: Comparison between the chl-roGFP2 redox state in young and old leaves under steady-state conditions.** A, Comparison between chl-roGFP2 $R_{405/465}$ values in young leaves (YL) and old leaves (OL) during rest, under fully oxidized (1000mM $H_2O_2$) and fully reduced (100mM DTT) conditions. The $R_{405/465}$ values are presented as a box plot (n=3). B, Whole-plant imaging of plants grown under constant light conditions (300-µmol photons $m^{-2}$ $s^{-1}$), highlighting the spatial distribution of redox changes between young, and old leaves.
